# Stoichiometric quantification of spatially dense assemblies with qPAINT

**DOI:** 10.1101/525345

**Authors:** Matthew AB Baker, Daniel J Nieves, Geva Hilzenrat, Jonathan F Berengut, Katharina Gaus, Lawrence K Lee

## Abstract

Quantitative PAINT (qPAINT) is a useful method for counting well-separated molecules within nanoscale assemblies. But whether cross-reactivity in densely-packed arrangements perturbs measurements is unknown. Here we establish that qPAINT measurements are robust even when target molecules are separated by as little as 3 nm, sufficiently close that single-stranded DNA binding sites can interact.

## 1. Introduction

Super-resolution microscopy enables the distinct observation of molecules that are separated by a distance that is less than the diffraction limit of light. In biological systems, it has been used to elucidate the nanoscale distribution and organization of many different fluorescently labelled molecules in unprecedented detail (*1*-*3*). Super-resolution imaging can be achieved through single molecule localization microscopy (SMLM) where molecules are imaged independently by separating their fluorescence emission in time, then fit to a point spread function (PSF) to determine their locations (*4*). Traditionally, SMLM was implemented by two approaches that utilize the stochastic photoswitching of an optically resolvable subset of fluorophores: photo-activatable and photo-convertible localization microscopy (PALM) (*5*) and stochastic optical reconstruction microscopy (STORM) (*6*). Both have proven to be to be exceptionally useful for imaging of many structures, from cell surface receptors to cytoskeletal components (*7, 8*). Since PALM and STORM rely on fixed fluorescent labels, they are subject to the photo-physical properties of the fluorescent label. This includes a limit to the number of emitted photons before irreversible photobleaching occurs (*9*), which can lead to poor signal (*10*).

In an alternative method known as point accumulation for imaging nanoscale topography (PAINT), SMLM can be achieved by imaging target sites *via* the stochastic and reversible binding of rapidly diffusing fluorescently labelled ligands in solution (*11*-*13*). Jungmann *et al.* has implemented the PAINT method with a DNA-based system (DNA-PAINT) which utilises the programmable, specific binding of a DNA-labelled ‘imager’ fluorophore to a complementary ‘docking’ strand (*14*-*16*). PAINT involves the continuous exchange of fluorescent ligands and data can in principle be accumulated indefinitely, while binding sites remain intact (*17*). The same group recently extended the DNA-PAINT method by demonstrating that the time interval between binding events, or dark time, is inversely proportional to the number of binding sites (*15*). Consequently, the number of docking strands could be quantified using a method known as quantitative PAINT (qPAINT). By normalizing to the dark time of a single binding site, qPAINT was used to count up to 12 docking strands separated by 20 nm on a DNA origami structure (*15*).

qPAINT has already been utilized to tackle one of the most difficult problems in biology: quantifying the stoichiometries of clustered proteins in cells. For example, qPAINT was initially used to observe the number of CAZ units bound to Bruchpilot proteins, with approximately 140 Bruchpilot proteins measured per CAZ unit (*15*). More recently, qPAINT was used to investigate the distribution of signaling ryanodine receptors (RyRs) within the membrane of cardiac myocytes (*18*). Jayasinghe *et al.* were able to resolve single RyRs within the plasma membrane, as they were separated by 20-40 nm, and confirmed this with qPAINT quantification. However, the dimensions of some proteins and their intermolecular distances in assemblies are comparable to the length of the 10 nucleotide docking strands typically used in DNA-PAINT experiments (∼3 nm) (*19, 20*). Thus, it is conceivable that when docking strands are so densely packed, cross-reactivity between imager strands will alter binding kinetics(*21, 22*), significantly confounding the mean dark times measured from qPAINT experiments. We, therefore, sought to address the question of whether qPAINT is a robust method for counting integer stoichiometries when docking strands are separated by small distances. We used a new DNA nanotube structure (*23*), which allowed us to compare dark times between four identical DNA-PAINT sites that were separated by ∼15 nm and as little as 3 nm.

## 2. Methods and Materials

### 2.1. DNA Origami Synthesis

52-helix pleated DNA origami nanotubes were synthesised as previously described (*23*). Four staple extensions (5’-CGACACCTTAAAGCCGCCGC-3’) that were complementary to 20 nt Biotinylated DNA strands (5’-BIOTIN/GCGGCGGCTTTAAGGTGTCG-3) were incorporated at one end of the nanotubes to immobilise the nanotubes on streptavidin-coated surfaces. A further four 20 nucleotide staple extensions were placed on the same, lower, surface-attached end of the nanotubes to bind to complementary fluorescent oligomers labelled with ATTO565 or Alexa488 to locate and identify DNA nanotubes (5’-CGCCCGCTGAAAAAGCTGCG-3’ and 5’-ATTO565N/CGCAGCTTTTTCAGCGGGCG-3’; 5’-CGCCCGCAAGTCTCACCGCG-3’ and 5’-Alexa488N/CGCGGTGAGACTTGCGGGCG-3’ respectively). Docking strands consisted of 10 nucleotide single strand sequences (docking sequence: TCCTCAATTA). These were attached to the pleated nanotube by extending the 3’ end of DNA staples positioned at the opposite end of the pleated nanotube. These docking strands were located on the inner DNA helices and transiently interacted with imager strands (TAATTGAGGA-Alexa647N) that were complementary to the docking sequence (Fig. 1). An antiparallel-parallel-antiparallel (APA) staple was incorporated into the 4SEQ nanotubes. This created DNA staple crossovers between the inner DNA helices of the nanotube resulting in a more rigid structure where expansion from electrostatic repulsion between DNA helices was prevented (*23*). This allowed parallel docking strands to be closely spaced (∼3 nm apart).

**Fig. 1.**
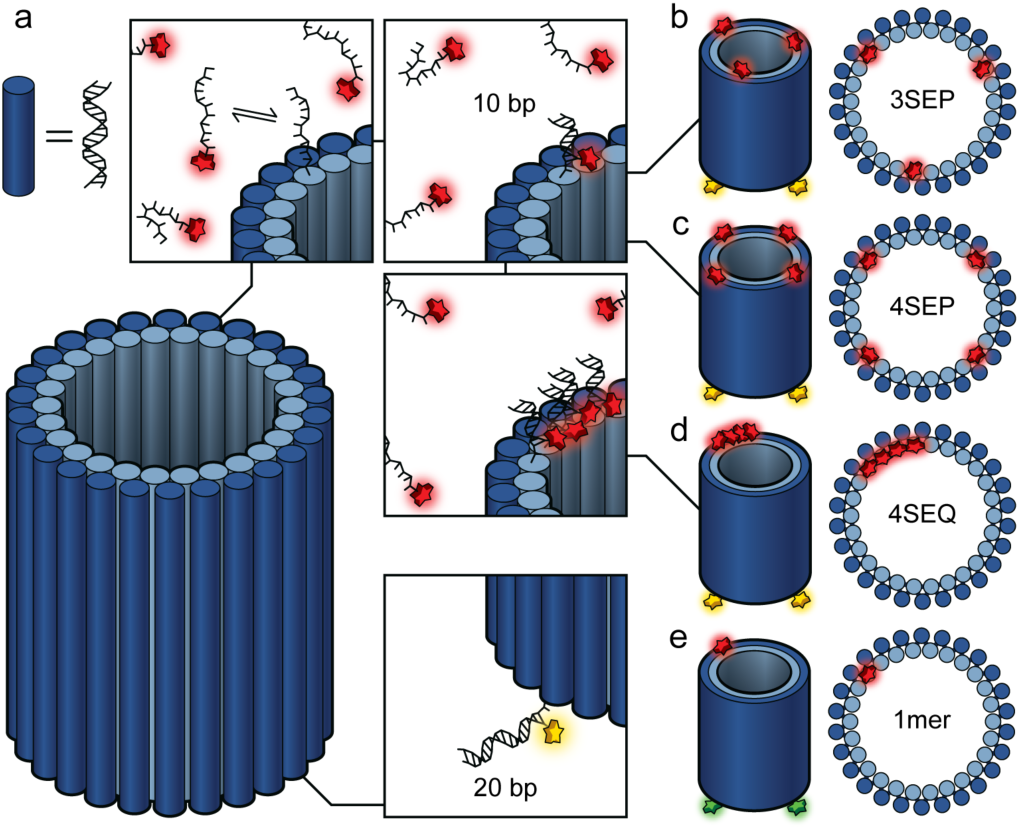
Schematic of DNA nanotube. (a) DNA-PAINT docking strands are incorporated as 3’-DNA staple extensions at the top of the 52-helix nanotube, with 10 nt sequence complementary to a DNA-PAINT imager strand labelled with Alexa647(red). Imager strands are transiently bound and exchange in solution. Three types of multimeric nanotube (b) 3SEP labelled at the top with 3 evenly spaced PAINT docker sites, (c) 4SEP labelled at the top with 4 PAINT docker sites separated by ∼15 nm, and (d) 4SEQ labelled at the top with 4 closely packed PAINT docker sites (∼3 nm separation). The base of each multimeric nanotube has four 20 nt staple extensions which are complementary to oligonucleotides labelled with ATTO565 (yellow) for locating and identifying the nanotubes (e) Single-site origami nanotube (1mer) used for qPAINT calibration. The base of the 1mer has four 20 nt staple extensions complementary to oligonucleotides labelled with Alexa488 (green).

### 2.2. Sample Preparation for Microscopy

Plasma-cleaned coverslips were mounted onto ethanol-cleaned microscope slide with double-sided tape to create a capillary channel. To minimise non-specific binding to the surface, 1 mg/mL biotin labelled albumin (A8549 Sigma Aldrich) in Buffer A+ (10 mM Tris-HCl, 100 mM NaCl, 0.05% Tween 20, pH 8.0) was flowed into the chamber by capillarity and incubated for 2 mins. Unbound albumin was then washed away by Buffer A+. Then 0.5 mg/mL streptavidin in Buffer A+ was flowed into the chamber and incubated for a further 5 mins. Excess streptavidin was washed away using Buffer A+, and 200 pM of 3/4mer DNA origami labelled with ATTO565 mixed with 200 pM monomer labelled with Alexa488 in Buffer B+ (5 mM Tris-HCl, 10 mM MgCl2, 1 Mm EDTA, pH 8.0, Tween 20 0.05% vol/vol) was flowed into the chamber and incubated for 5 mins. Unbound origami was washed away with Buffer B+. A 1 in 500 dilution from a stock of gold nanorods 25 × 65 nm (cat. A12-25-650, Nanopartz, USA) were flowed into the chamber as fiduciary markers for subsequent drift correction, incubated for 10 minutes, then washed out with Buffer B+. Freshly prepared Alexa647-DNA imaging strands (10 nM diluted from 1 μM stock) in Buffer B+ were flowed into the chamber and the chamber was sealed at both ends using VALAP.

### 2.3. Imaging

Images were acquired using a Zeiss Elyra P.1 microscope using total internal reflection fluorescence (TIRF) with a Zeiss Alpha Plan Apochromat 100x NA 1.46 oil objective. First, to locate DNA origami structures, a time series of images of the 3/4mer ATTO565 labelled origami nanotubes was taken (0.5 kW/cm^2^, 100 ms, 500 frames). This was then repeated for the Alexa488 labelled origami nanotubes with a single docking strand for 500 frames with an integration time of 100 ms/frame under 0.004 kW/cm^2^ 488 nm laser illumination with a TIRF angle of 66.90°. For DNA-PAINT imaging, 642 nm (0.075 kW/cm^2^) laser illumination was used with an integration time of 300 ms for 50,000 frames with a TIRF angle of 66.90°. All images were 512×512 pixels with a pixel size of 97 nm.

### 2.4. Image Processing

DNA-PAINT images were processed using Zeiss Zen Black software. DNA-PAINT localisation data (Alexa647) was drift corrected using the point patterns generated from the localisation of gold nanorod fiducials within the field of view using the Zeiss Zen Black software drift correction and the resulting corrected localisation table was used for further analysis. After Gaussian filtering, PSFs from blinking events within single image frames were firstly selected according to the pixel radius of the circular features present in each frame using a peak mask radius of 7 pixels. Secondly, the PSFs were further filtered according to their signal to noise ratio when I-M > S*SNR, where I is the PSF intensity, M is the mean image intensity, S is the standard deviation of the image intensity, and SNR is the minimum signal to noise ratio threshold, here a value of 6. Each PSF that passed these criteria was then fitted to a two-dimensional Gaussian distribution to find the centre and the localisation precision. Localisation precision was calculated by published methods^3^. This was repeated for all frames in the stack and the x,y-coordinates were recorded for further processing (Fig. 2a). To determine the position of the DNA origami nanotubes a mean intensity image was calculated from 500 frame image stack of the tightly bound fluorophores (ATTO488 for monomer and ATTO565 for trimer/tetramer, illuminated using either 488 nm or 561 nm, respectively) prior to DNA-PAINT imaging. The single mean intensity image was then processed as above using 2D Gaussian fitting to find the centre of the permanently labelled DNA origami nanotubes within the field of view. These coordinates were recorded for the generation of the masks for further analysis.

**Figure 2:**
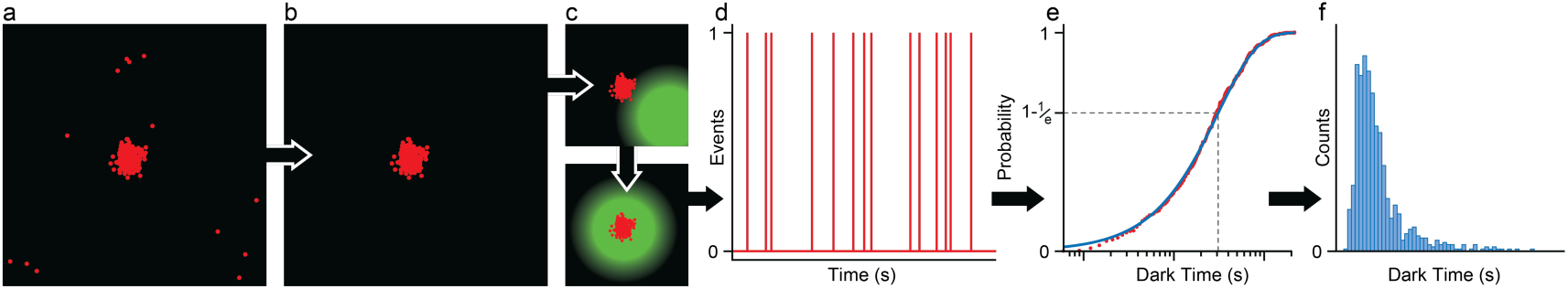
qPAINT analysis pipeline for DNA origami nanotubes. (a) The centres of origami-bound imager strands for a representative 4SEP nanotube are determined by fitting with a 2D Gaussian point spread function (PSF), which can then be rendered as a point pattern (red). (b) The point pattern data is then segmented using DBSCAN(*24*). and clusters that do not satisfy the DBSCAN parameters constitute background detections and are removed. (c) The position of the centre of each labelled nanotube is determined by 2D Gaussian fitting to the fluorophore at the base of the nanotube (ATTO565 or Alexa488 respectively). These are then aligned with the cluster of points from bound imager strands (bottom. (d) The arrival of each imager and the dwell times between events can be plotted vs time (magenta, bound = 1, unbound = 0). (f) Dark times are defined as the time between binding events and are plotted as a plotted as a cumulative distribution function (CDF). This CDF is fitted with a single exponential to extract of the mean dark time for each nanotube corresponding to (1-1/e) of the CDF. (f) The mean dark times from each individual CDF are then aggregated into a single histogram.

### 2.5. Cluster segmentation and masking

The x,y coordinates of imager binding events were clustered using the DBSCAN algorithm(*24*) with a minimum point per cluster (minPts) = 30 and a radius of search around single points (ε) = 15 nm (Fig. 2b). The indices of DNA-PAINT localisations within clusters were recorded and used to select data points corresponding to the same cluster from the localisation table for generation of time traces in later analysis. The coordinates from the centres of labelled origamis within the mean mask images were convolved with a 2D Gaussian and aligned with the DNA-PAINT localisation data, also convolved with a 2D Gaussian, using a lateral transformation, calculated from spatial correlation of the two images (Fig. 2c). This allowed separation of trimer/tetramer and monomer clusters automatically within the same sample. Clusters falling within the masked region (circle of radius 50 nm) for either mask were assigned to a single DNA nanotube origami and then analysed for the frequency of events from binding/unbinding of detected DNA imaging strands.

### 2.6. Analysis of Dark Times

First, time traces for events in each cluster over time were generated from the first detected binding event until the last (Fig. 2d). A histogram of the dark times (time between binding events), for that cluster was then used to plot the exponential cumulative density function (CDF) for that cluster (Fig. 2e). Dark times corresponding to the length of one frame (here, 300 ms) were removed, as these events arise from blinking of the bound imaging strand. The exponential CDF of the dark times was fitted to a single exponential, and the mean dark time of the CDF extracted (mean of exponential CDF = 1-1/*e*).

### 2.7. Normalisation of Dark Time Histograms

Dark times were calculated as above and aggregated into histograms corresponding to each subpopulation with a bin width of 5 s (Fig. 2f). These histograms were fitted with normal distributions to determine the mean dark time and standard variation for each subpopulation. In order to account for variations in imager strand concentrations between different experiments, we normalised dark time measurements between two samples via the ratio of the mean monomer dark times in each sample. For example, the mean monomer dark time in each measurement was compared to the mean monomer dark time from the first monomer/4SEP sample (chosen as the normalisation baseline at ∼100 s) to calculate a multiplier that was then used to normalise the 3mer/4mer dark times. Following normalisation, dark time measurements from identical samples were pooled and then plotted with a fitted normal distribution (Fig. 3). Error was propagated by calculating the 95% confidence interval (CI) for the centre of the Gaussian fit across each individual measurement before normalisation, and also by calculating the 95% CI for the final pooled, normalised fit. Total error in the final mean measurement, Δτ_d_, was then determined by summing these absolute errors in quadrature. The mean dark time was then reported as τ_d_ ± Δτ_d_ for 3SEP, 4SEP and 4SEQ, where ± Δτd represents the Standard Error in the Mean (SEM)

**Figure 3:**
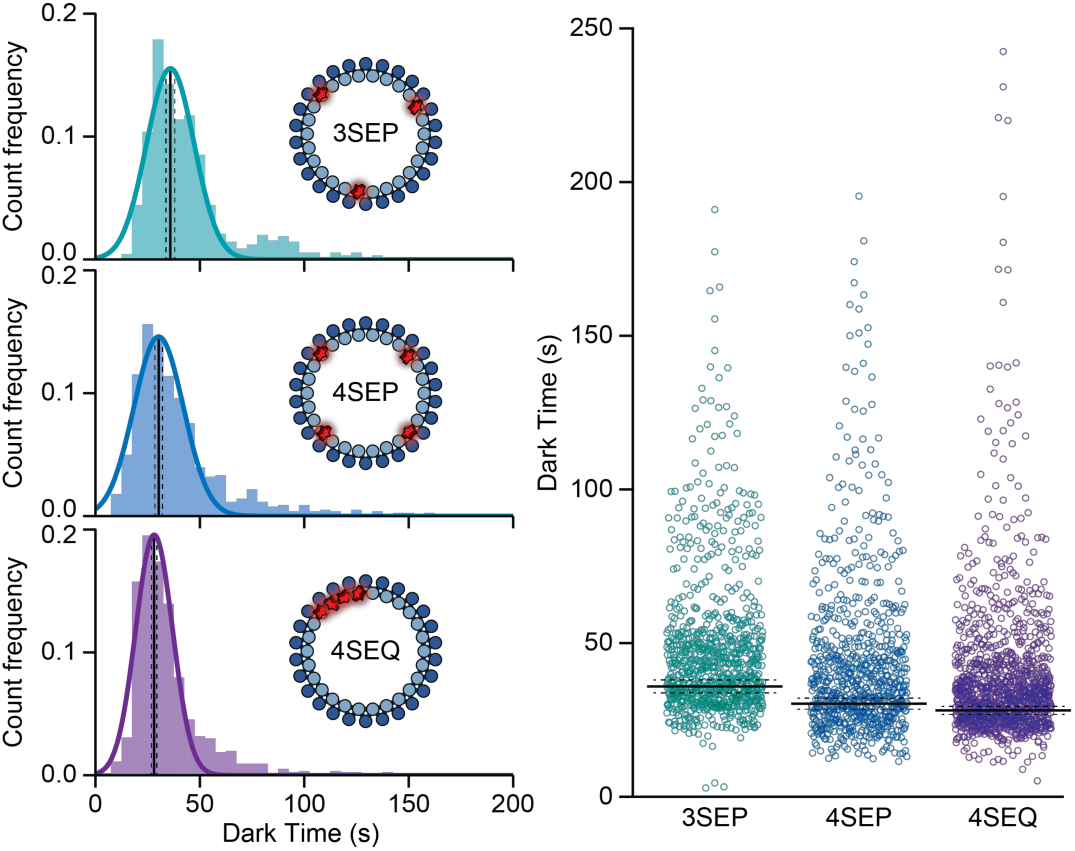
Dark time comparisons between 3SEP, 4SEP and 4SEQ. (left) Histograms of mean dark time for each individual DNA origami nanotube fit by a single Gaussian and showing the centre of the fit (solid black line) and the standard error associated with that fit (dashed lines) (top: 3SEP; middle: 4SEP; bottom: 4SEQ). (right) Spread plot of mean dark time showing that the mean dark time of 4SEP and 4SEQ overlap within error whilst 3SEP is distinguishable.

## 3. Results and Discussion

qPAINT experiments were performed on a 52-helix pleated DNA origami nanotube (*23*) with parallel docking strands arranged at one end of the nanotube (Fig. 1). The pleated nanotube design allowed for parallel docking sites to be as close as 3 nm (Fig.1a). We measured the mean dark times associated with binding of Alexa647 labelled imager strands on DNA origami nanotubes with different docking strand configurations with total internal reflection fluorescence (TIRF) microscopy. Nanotube configurations consisted of three (3SEP; Fig. 2b) and four (4SEP; Fig. 2c) docking strands separated by ∼ 20 nm as well as on a third DNA origami nanotube with four docking strands placed on sequential helices separated by 3 nm (4SEQ; Fig 2d). Binding events were localized to individual origami structures and dark times normalized by comparison with nanotubes with a single binding docking strand (1mer; Fig. 1e). These were mixed with each sample and imaged simultaneously to control for variations in imager concentration. To distinguish between 1mer control nanotubes and those for qPAINT experiments all nanotubes possessed another tightly bound fluorophore at their base (ATTO565 and Alexa488 for target and 1mer respectively; Fig. 1).

The positions of bound imager strands within the DNA-PAINT images were determined by 2D Gaussian fitting of the point spread functions from individual binding events to yield single molecule localisation data. These consisted of clusters of points corresponding to the specific binding of imager strands to docking strands, as well as isolated points corresponding to non-specific background interactions of imager strands to the BSA-blocked surface (Fig. 2a). These background points were automatically removed by processing images with a propagative density based clustering algorithm, DBSCAN(*24*) (Fig. 2b). Clusters of points in single molecule localisation data could be aligned with the centre of PSFs from images acquired of fluorophores bound tightly to the base of the DNA origami nanotubes (Fig. 2c). This confirmed that data corresponded to interactions between imager strands located on the DNA nanotube. For each DNA nanotube, binding events were separated temporally to determine dark times (τ_d_) (Fig. 2d), which were plotted as a cumulative density function (CDF). These were consistent with a single exponential from which the mean dark time (1-1/e) for each structure was determined (Fig. 2e). The mean dark time for each cluster was then aggregated into a histogram, from which the overall mean dark time for that population of origami nanotube was extracted (Fig. 2f).

Mean dark times differed between nanotubes with three docking sites (35.9 ± 2.1 s) and four docking sites but not between 4SEP (30.3 ± 1.8 s) and 4SEQ (28.1 ± 1.3 s), where docking strands are separated by only 3 nm, sufficiently close to cross-react with neighboring strands (Fig. 3 and Fig. S1). These data demonstrate that closely spaced docking strands do not significantly affect stoichiometric measurements with qPAINT. Moreover, dark times were inversely proportional to the number of docking sites (0.84 ± 0.07 and 0.78 +/- 0.06 for 4SEP and 4SEQ respectively) and similar to the ideal ratio between three and four docking sites (0.75). This confirms the robustness of our experimental system and analysis pipeline to quantify integer stoichiometries of docking strands.

## 4. Conclusions

We have demonstrated that closely spaced docking strands do not affect stoichiometric measurements with qPAINT. We thus establish the suitability of qPAINT for use in determining stoichiometries in samples with densely packed spatial arrangements such as in biological macromolecular assemblies.

## Author Contributions

M.A.B.B. conceived the project, synthesised DNA origami, collected and analysed data and wrote the manuscript, D.J.N. collected and analysed data and wrote the manuscript, G.H. analysed data, J.F.B. designed DNA origami and wrote the manuscript, K.G. wrote the manuscript, L.K.L. conceived and supervised the project, and wrote the manuscript.

## Supplementary Figure

**Figure S1.**
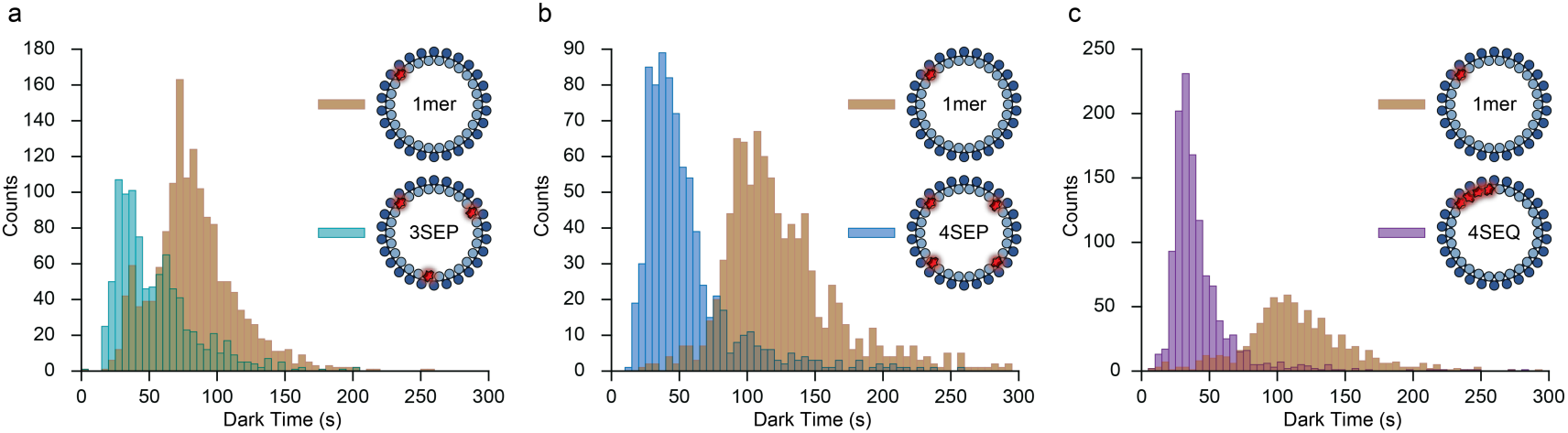
Histograms of dark times prior to normalisation. Histograms of dark times for all DNA origami nanotubes imaged in samples containing (a) 3SEP, (b) 4SEP and (c) 4SEQ. Dark times for nanotubes with a single docking site (1mer) for normalization are shown in orange.

